# Exploring the Aβ Plaque Microenvironment in Alzheimer’s Disease Mice by Multimodal Lipid-Protein-Histology Imaging on a Benchtop Mass Spectrometer

**DOI:** 10.1101/2024.12.04.626756

**Authors:** Elisabeth Müller, Thomas Enzlein, Dagmar Niemeyer, Livia von Ammon, Katherine Stumpo, Knut Biber, Corinna Klein, Carsten Hopf

## Abstract

Amyloid-β (Aβ) plaque deposits in the brain are a hallmark of Alzheimer’s disease (AD) neuropathology. Plaques consist of complex mixtures of peptides like Aβ_1-42_, characteristic lipids such as gangliosides, and they are targeted by reactive microglia and astrocytes. In pharmaceutical research and development it is therefore a formidable challenge to contextualize different biomolecular classes and cell types of the Aβ plaque microenvironment in a coherent experimental workflow on a single tissue section and on a benchtop imaging reader. Here, we developed such a workflow that combines lipid MALDI mass spectrometry imaging using a vacuum-stable matrix with histopathology stains and with MALDI HiPLEX-immunohistochemistry of plaques and multiple protein markers on a bench-top imaging mass spectrometer. The three data layers consisting of lipid, protein markers, and histology can be co-registered and evaluated together. Multimodal data analysis suggests extensive co-localization of Aβ plaques with the peptide precursor protein, with a defined subset of lipids and with reactive glia cells on a single brain section of APP/PS1 AD model mice. Plaque-associated lipids like ganglioside GM2 and phosphatidylinositol PI38:4 isoforms were readily identified using the tandem MS-capabilities of the mass spectrometer. Taken together, our data suggests that complex pathology involving multiple lipids, proteins and cell types can be interrogated by this spatial multiomics workflow on an easy-to-use benchtop mass spectrometer.

## 1. Introduction

Alzheimer’s disease (AD), a progressive neurodegenerative disorder, is the leading cause of dementia in the elderly population that affects over 50 million people globally. Numbers are expected to rise, as populations age [1,2]. AD is characterized by cognitive decline, memory loss, and behavioral changes. Key pathological features include extracellular amyloid-beta (Aβ) plaques, intracellular neurofibrillary tangles (NFTs) of hyperphosphorylated tau, and more recently, neuroinflammation driven by reactive microglia and astrocytes as a potential third AD hallmark [3,4]. Aβ plaques in AD primarily consist of Aβ peptides, formed by abnormal cleavage of amyloid precursor protein (APP) by β- and γ-secretase enzymes [5]. The cellular and molecular microenvironment around Aβ plaques is complex, and it influences disease progression. Immune cells like microglia and astrocytes cluster around plaques, initially clearing Aβ, but eventually becoming dysfunctional with chronic responses, e.g., releasing pro-inflammatory cytokines [6,7]. Altered lipid metabolism, especially of cholesterol and gangliosides, in the vicinity of plaques may contribute to further disease progression [8,9]. Several AD mouse models have been genetically engineered to replicate key AD features, such as Aβ plaques, tau pathology, and in some cases cognitive decline. Transgenic models, like APP/PS1 mice, overexpressing human genes linked to early-onset familial AD (e.g., mutant APP and presenilin 1 (PS1)), are used in pharmaceutical R&D to study Aβ pathology and plaque formation [10]. AD current treatments only provide symptomatic relief or modify the disease in minor ways [11]. However, advances in genomics, biomarkers, and imaging have offered new insights into potential therapeutic targets, thus fueling optimism for new drug classes [12,13]. New technologies for pharmaceutical inquiry have become available as well.

Matrix-Assisted Laser Desorption/Ionization Mass Spectrometry Imaging (MALDI MSI) is an advanced technique used to visualize the spatial distribution of different biomolecular classes in tissue sections coated with a chemical matrix. MALDI MSI enables the creation of molecular maps of the tissue, allowing visualization without the need for labels or dyes [14]. It detects a wide range of biomolecules, from small metabolites to proteins, depending on the matrix and conditions [14]. It can offer high spatial resolution, typically from tens to hundreds of micrometers [15]. Technological advances are now enabling studies even at cellular levels [16]. MALDI MSI has been widely used in AD research to investigate the brain’s molecular landscape [17–21]. Multiple studies focused on mapping the spatial distribution of Aβ peptides and associated lipids from brain tissue sections, thereby enabling studies of plaque location, composition and differentiation between various Aβ isoforms, [17,19]. MALDI MSI has been used to visualize Aβ plaques in transgenic AD mouse models and human AD patient samples, revealing that the distribution of Aβ species correlates with other pathological features such as microglia cells as well as lipid species [18,21–23].

Biochemical and clinical studies indicate that, alongside Aβ and tau pathology, alterations in neuronal lipid metabolism are involved in AD disease progression [24]. The ε4 allele of the apolipoprotein E encoding gene (APOE) is one of the most important genetic risk factor in AD [25]. In recent years, genome-wide association studies have identified more lipid-associated genes as risk factors, like the lipid transporter TREM2, highlighting the potential of lipid metabolism as new therapeutic target [26]. Current research focuses on identifying region-specific lipid changes in AD brains compared to healthy controls, particularly in areas like the hippocampus and cortex which are affected by plaque depositions in the progression of the disease [20,23]. Accumulation of distinct lipid species, such as glycerophospholipids and sphingolipids like gangliosides were identified in the vicinity of Aβ plaques in AD mouse models and human AD patient samples using MSI [18,21,23]. Some lipids, such as gangliosides, interact with Aβ peptides and promote their aggregation into plaques [9]. The metabolism of other lipid classes, such as derivatives of phosphatidylinositol, seem to be disrupted by Aβ oligomers [27]. The increase of ceramides can be related to the observed decrease of sphingomyelin (namely sulfatides) in AD mouse brain samples, where sulfatides might be degraded during the progression of AD [28].

To comprehensively assess the complexity of AD, it is fundamentally important to consider as many classes of biomolecules as possible [29]. For years, large molecules such as amyloid beta peptides and phosphorylated tau have gained most of the attention in AD research [30]. Recently, there is a growing interest in the metabolic activities of cells in the plaque microenvironment [4]. Ideally, the analysis of lipids, peptides, and proteins would be accomplished on a benchtop imaging reader from one single tissue sample. However, while the analysis of lipids is very common using MALDI MSI, the detection of large molecules, such as intact proteins, is more challenging [31]. Strategies like on-tissue digestion of the proteins result in very complex data sets with high background that are limited to highly abundant peptides and provide very low sequence coverage of proteins [32,33]. Mostly, proteins of interest are investigated by immunohistochemistry (IHC), with one or even multiple (high-plex) antibody labels to the tissue sample [34,35]. Consequently, lipid imaging and protein IHC detection from one single tissue slide is very challenging, and experiments are typically performed on two consecutive sections using high-end mass spectrometers instead of easy-to-use benchtop instruments [36].

Here, we demonstrate the feasibility of multiomic and multimodal MALDI MSI of lipids, protein markers, and H&E histology, all on the same tissue section and, on a benchtop MALDI imaging instrument. Using MALDI HiPLEX-IHC technology [37] to visualize amyloid-β plaques and known cell/protein markers of their proinflammatory cellular microenvironment, we contextualize lipids as key components of the plaque microenvironment in a mouse model of AD. Finally, we demonstrate the ability to elucidate lipid structures directly on tissue with MS2 fragmentation analysis, all on the same instrument..

## 2. Results

### 2.1. Workflow for multimodal lipid MSI, MALDI HiPLEX-immunohistochemistry (IHC) and H&E histology on a benchtop mass spectrometer

In this study, we sought to develop a new workflow for multimodal lipid MSI and MALDI HiPLEX-immunohistochemistry (IHC) on the same tissue section. Our goal was to do this on a newly available benchtop MALDI-TOF/TOF mass spectrometer. To this end, a single cryosection of fresh-frozen APP/PS1 mouse brain or a wild-type control mouse were spray-coated with the vacuum-stable MALDI matrix DMNB-2,5-DHAP **[38]** for untargeted lipid MS imaging for *m/z* 400−1600 in negative ionization mode. Subsequently, MALDI matrix was removed for further processing. For MALDI HiPLEX-IHC on the same sample section, the tissue was fixed in paraformaldehyde and washed to remove lipids and metabolites. The brain section was then incubated with multiple antibodies simultaneously. Each antibody is conjugated to a distinct photo-cleavable peptide mass tag (PCMT) as a unique reporter for a single protein marker. After staining and UV-cleavage of mass tags from the antibodies, the sections were spray-coated with the MALDI matrix CHCA, followed by a second MS imaging run in positive ionization mode to record the PCMT reporters. This allowed us to acquire targeted peptide mass tag data for *m/z* 600−4000 (**Figure 1a**) in addition to the lipid MSI data.

**Figure 1.**
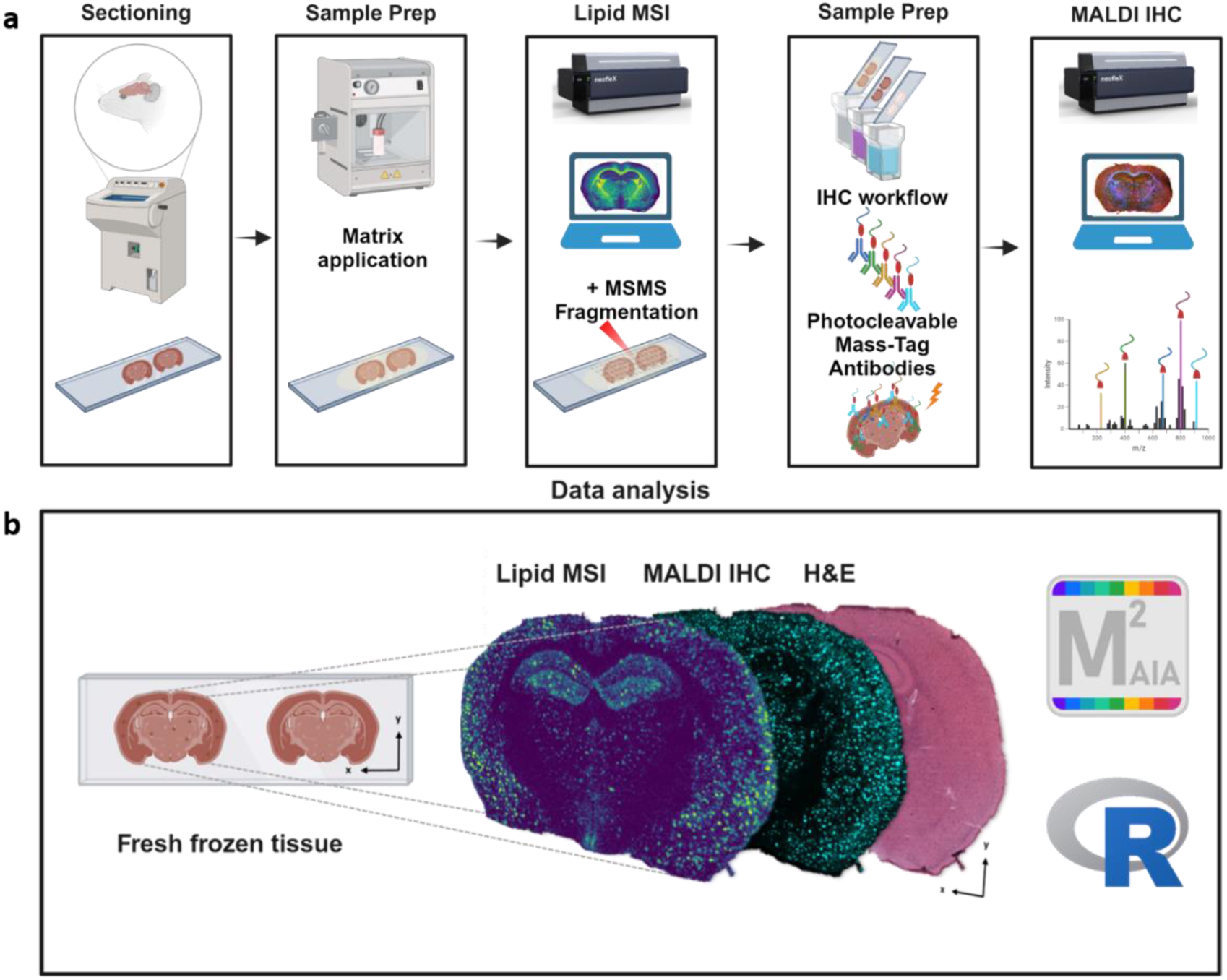
Benchtop mass spectrometry workflow for multimodal MALDI lipid imaging, MALDI HiPLEX-immunohistochemistry (IHC) and hematoxylin & eosin (H&E) histology of the Aβ plaque microenvironment in Alzheimer’s disease (AD) model mice. a, Fresh-frozen brains from the AD mouse model APP/PS1 were cryosectioned, mounted on conductive ITO glass slides and spray-coated with the vacuum-stable MALDI matrix DMNB-2,5-DHAP [38]. Lipid MSI data was acquired on a neofleX MALDI-TOF benchtop mass spectrometer from full brain sections in negative ionization mode. MALDI matrix was then removed. For MALDI HiPLEX-IHC, the brain section was incubated with a mix of antibodies, each conjugated to a distinct PCMT as a reporter for a single antigen/protein marker. A second “protein” MSI run imaged the reporter mass tags on the same tissue section in positive ionization mode. The neofleX TOF/TOF capability was used to obtain MS2 fragment spectra for a predefined list of *m/z* features, namely lipids, for structure elucidation on a consecutive tissue section. b, Individual ion images of PCMTs were visualized in SCiLS Scope software, and co-registration of lipid, protein, and H&E modality layers was performed in M^2^aia software [18,39].

For correlative investigation of multimodal data, the two MSI data sets and the H&E image obtained from an adjacent slide were co-registered using in-house M^2^aia software (**[39,40]**). This made visualization of all the three layers (H&E, lipids, and protein HiPLEX-IHC) possible (**Figure 1b**). For data analysis, we reversed the order used for data acquisition (lipid first, then HiPLEX-IHC) and focus on spatial analysis of protein markers first.

### 2.2. MALDI HiPLEX-IHC staining reveals AD biomarker co-localization in the microenvironment of Aβ_1-42_- positive plaques in APP/PS1 mouse brain

The AD research focus has expanded beyond Aβ plaques and hyper-phosphorylated tau proteins. It now includes additional molecular characteristics in close association with Aβ deposits. The testable hypothesis is that Aβ peptide deposition, tau pathology, and reactive immune cells like microglia together drive neurodegeneration in AD and ultimately dementia [4]. This shows the importance of uncovering the microenvironment of Aβ plaques to fully understand the progression of the disease. Using the new workflow on a benchtop mass spectrometer, we investigated the surroundings of Aβ_1-42_- positive plaques using multiple PCMT-labeled antibodies to localize multiple neurology markers in one AD mouse brain section. An H&E image provided tissue morphology information, including the cortex and hippocampal regions as landmarks within the mouse brain tissue sample (**Figure 2a**) [41]. Structural markers such as NeuN included in the HiPLEX antibody panel supported navigation of relevant brain areas, e.g., the pyramidal cell layer of dentate gyrus in the hippocampus (**Figure 2b**). We detected many dense Aβ plaques using an antibody against Aβ_1-42_ (ion image of its reporter mass tag at *m/z* 1771.5; **Figure 2c**) in various brain areas including the cerebral cortex and hippocampus, in accordance with other studies of this AD mouse model using other imaging technologies [10]. No Aβ_1-42_-positive regions were observed in WT mouse sample **(Supplementary Figure 1)**. Microglia were detected with anti-Iba-1 antibody (**Figure 2d**; single ion image of *m/z* 960.1 reporter tag) with high expression in hippocampus and cortex of 13-month-old APP/PS1, but not in 14-month-old wild type (WT) mice **(Supplement Figure 1)**. Furthermore, we investigated the astrogliosis marker of glial fibrillary acidic protein (GFAP, *m/z* 1011.9), which displayed an overall similar distribution pattern as Iba-1. Additionally, there was some visualization in the transitions of the hippocampal sub-regions (Figure 2e). For unknown reasons, this distribution was also detected in the WT sample **(Supplement Figure 1e)**. As a third marker we investigated localization of the amyloid precursor protein (APP, *m/z* 1723.6) that Aβ peptides are derived from. The spatial distribution of this marker also showed partial signal enrichment in cortex and hippocampus. Furthermore, we observed a presumably endogenous signal pattern of unknown origin in the outer hippocampus area (Figure 2f). This signal pattern was also observed in the wild type sample to some extent (Supplement Figure 1f).

**Figure 2.**
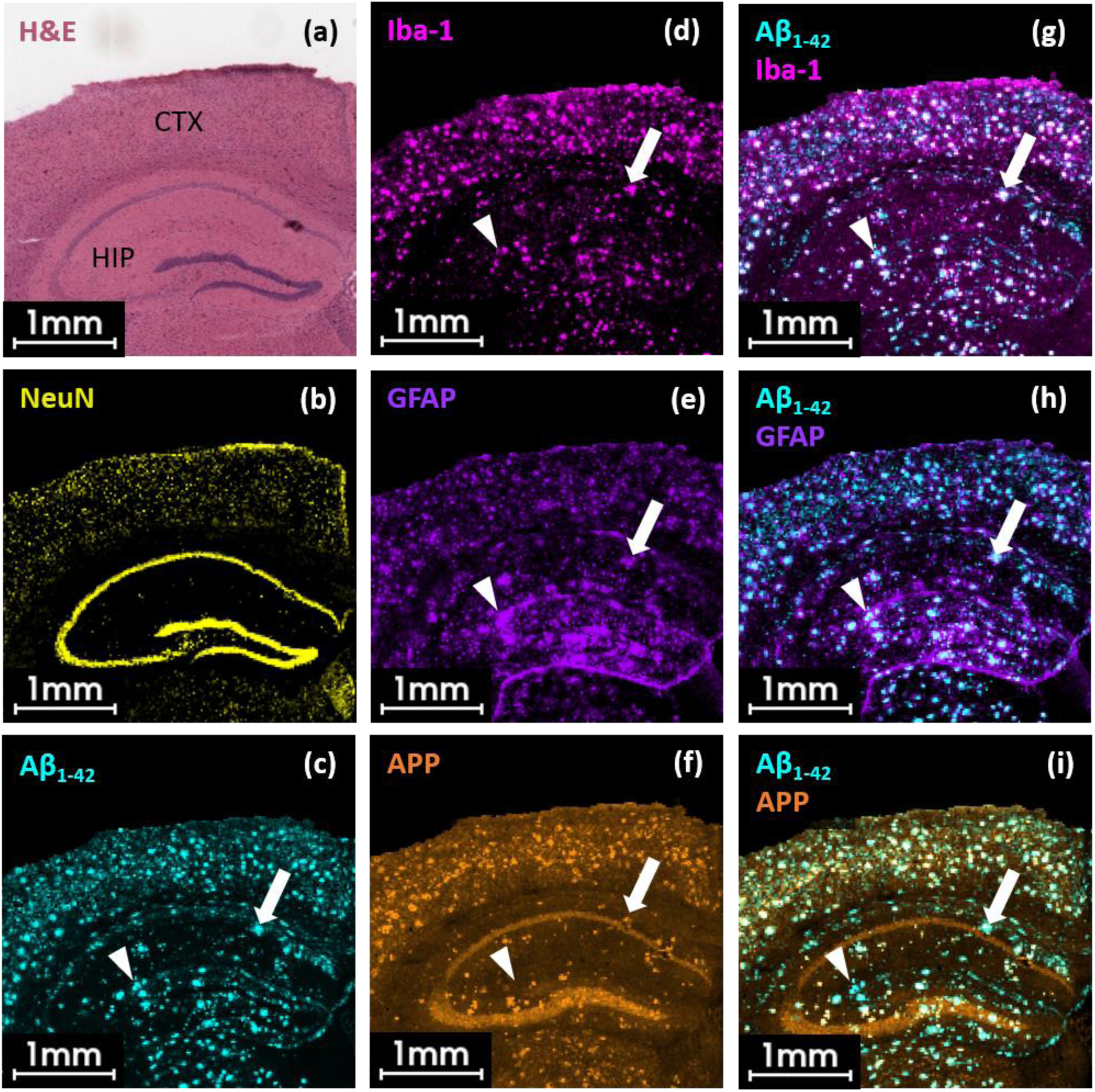
MALDI HiPLEX-IHC staining indicates extensive co-localization of Aβ_1-42_ plaques with microglia, activated astrocytes and the amyloid precursor protein APP in APP/PS1 mouse brain. MALDI HiPLEX-IHC images of five PCMTs indicating neuronal marker distribution in a coronal APP/PS1 mouse brain section. Mass reporters of the brain structural marker NeuN (Neuronal nuclei, *m/z* 1309.0 [M+H]^+^), the amyloid-β_1-42_ peptide (Aβ_1-42_, *m/z* 1771.5 [M+H]^+^), the microglia marker Iba-1 (*m/z* 960.1 [M+H]^+^), glial fibrillary acidic protein (GFAP, *m/z* 1011.9 [M+H]^+^), and the amyloid precursor protein (APP, *m/z* 1723.6 [M+H]^+^). Data was acquired in reflector positive mode at 20 *μ*m lateral step-size using a neofleX MALDI-TOF benchtop instrument. H&E staining (a) highlights distinct brain structures e.g., the hippocampus (HIP) and cortex (CTX). H&E staining is reflected by the structural marker NeuN (yellow, b) indicating regions with high density of neuronal cell bodies like the pyramidal neuron layer. Single ion image showing the spatial distribution of Aβ_1-42_-containing plaques (c), Iba-1-positive microglia (d), the astrogliosis marker GFAP (e), and the amyloid precursor protein (APP, f). Overlay of Aβ_1-42_ with Iba-1 (g), GFAP (h) and APP (i) show partial co-localization (white). Filled white arrow shows co-localization of Aβ_1-42_ with all other biomarkers. The filled white arrow head shows Aβ_1-42_ co-localization with Iba-1 and GFAP, but not for APP.

To further investigate the correlation between Aβ_1-42_-positive plaques and the three individual neurology markers, we superimposed each of these ion image with that of Aβ_1-42_ **(Figure 2g-i)**. The overlay of the Aβ_1-42_ signal (cyan) with Iba-1 (pink; microglia) revealed almost 100% co-localization of both in cortex and hippocampal regions (white, **Figure 2g**). White filled arrow and arrowhead indicate an example Aβ_1-42_ cluster (Figure 1c) in the hippocampus with co-localizing Iba-1 **(Figure 1d+g)**. We further explored ongoing astrogliosis in close vicinity to plaque material by overlaying the ion images of Aβ_1-42_ with the GFAP mass reporter. As for microglia, we detected multiple plaque sites co-localizing with GFAP (white filled arrow and filled arrowhead). However, we could detect high loads of GFAP-positive signals with huge clusters in hippocampus, whereas there seemed to be less signals co-localizing with Aβ_1-42_ in the cortex **(Figure 2h)**. Finally, in the APP/PS1 mouse, the overlay of Aβ_1-42_ (cyan) and APP (orange) suggested some degree of co-localization in cortex and hippocampus **(Figure 2i)**. For most Aβ_1-42_-positive stained regions, APP seemed to co-localize (white filled arrow). Nevertheless, some Aβ_1-42_-positive plaques seemed to not have APP in their surroundings (white filled arrowhead).

The intact protein imaging and co-localization of different neurological markers with Aβ_1-42_-positive plaque regions suggest a difference in the molecular composition of the regions of interest (ROIs). We therefore further investigated whether the composition around the Aβ plaques also differs at the lipid level.

### 2.3. Multimodal MSI reveals lipid composition of individual Aβ_1-42_-positive microglia regions in AD mouse brain tissue

Altered lipid homeostasis in the brain of AD patients became increasingly interesting, since multiple genome-wide associated studies (GWAS) found potential AD risk genes being associated with the lipid metabolism (reviewed in **[26]**).

We therefore extended the analysis of key protein markers in the Aβ plaque environment and combined it with the lipid data layer recorded earlier by untargeted MALDI lipid imaging. We investigated the co-localization of specific lipid species with Aβ_1-42_-positive plaque regions in the APP/PS1 mouse brain to identify potential lipid markers.

MALDI MSI of lipids was performed first on the tissue section that was followed by MALDI HiPLEX-IHC. Untargeted lipid data was acquired in a mass range of *m/z* 400-1600 using a vacuum-stable lipid matrix **[38]**. MALDI HiPLEX-IHC, MALDI MS lipid data and the respective H&E image of the tissue section were co-registered. ROIs were defined by Aβ_1-42_-reporter *m/z* 1771.5 **(Figure 3a (i))**. In total, fifty regions of Aβ_1-42_-positive areas in the cortex and hippocampus were manually selected from the MALDI HiPLEX-IHC data set **(Figure 3a (ii))**. Aβ_1-42_-containing ROIs can be displayed as an overlay on the H&E image for regional overview **(Figure 3a (iii)** and on single ion images of lipid features **(Figure 3b (i-viii))**.

**Figure 3.**
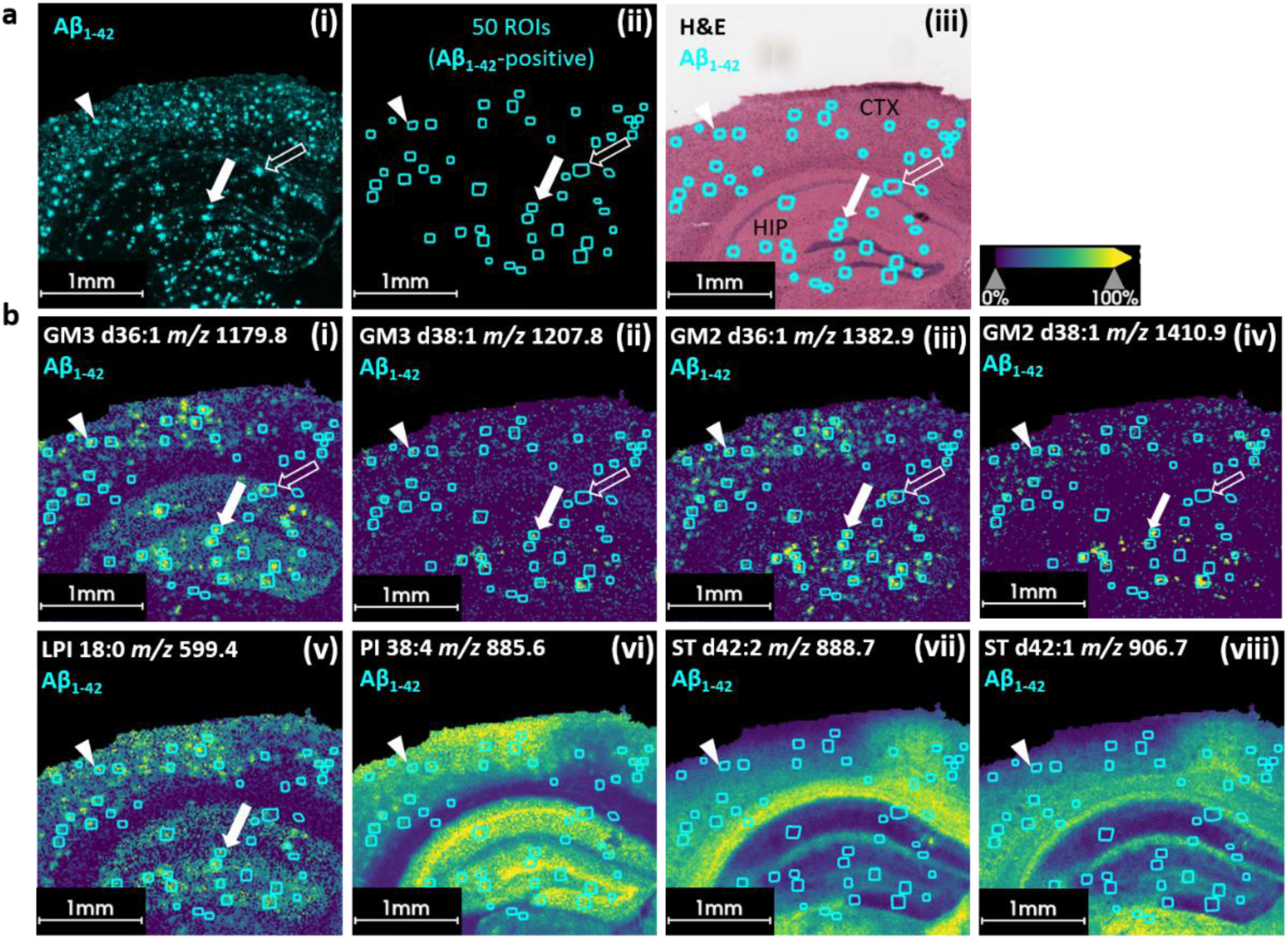
Multimodal MSI reveals lipid composition of individual Aβ_1-42_-positive plaque regions in AD mouse brain tissue. **a,** Ion images of the reporter mass for Aβ_1-42_ (*m/z* 1771.5 [M+H]^+^) showing general plaque distribution in cortex and hippocampus of APP/PS1 mouse brain (i). Fifty Aβ_1-42_ ROIs containing plaques were manually selected. (ii). Overlay of fifty Aβ_1-42_-positive ROIs with H&E image of the same tissue section (iii). **b,** Overlay of ion images of different lipid features in negative ionization mode with the fifty Aβ_1-42_-positive ROIs. (i-iv) Distribution of different GM3 and GM2 gangliosides in APP/PS1 mouse brain at plaque sites. White filled arrow indicates Aβ_1-42_-positive ROIs in (a) as well as lysophosphatidyl inositol (LPI 18:0, *m/z* 599.4, [M-H]^-^) accumulation in (b). (i-v). White unfilled arrows showing lipid deposition which is only true for GM3 d36:1 (*m/z* 1179.8, [M-H]^-^) and GM2 d36:1 (*m/z* 1382.9, [M-H]^-^), but not for GM3 d38:1 (*m/z* 1207.8, [M-H]^-^) and GM2 d38:1 (*m/z* 1410.9, [M-H]^-^), thus revealing possible differences in lipid composition surrounding plaques. (i-vi) White arrow heads showing accumulation of gangliosides and (lyso)-phosphatidylinositol as well as (vii-viii) depletion of two sulfatides (ST d42:2, *m/z* 888.7, [M-H]^-^ and ST d42:1, *m/z* 906.7, [M-H]^-^) in the plaque environment.

The visualization of single ion intensity distribution for individual lipid species showed a consistent deposit or depletion-like pattern throughout the cortical and hippocampal regions. We were able to show accurate correlation of lipid features such as sphingolipids with the Aβ_1-42_-positive plaque regions. The inspection of these ion images revealed plaque-associated accumulation of gangliosides (GM3 36:1, *m/z* 1179.8, GM3 38:1, *m/z* 1207.8, GM2 36:1, *m/z* 1382. 9, GM2 38:1, *m/z* 1410.9) in APP/PS1 brain. No substantial elevation of gangliosides was found in the control sample **(Supplement Figure 2)**. This is consistent with previous studies showing the accumulation of gangliosides at amyloid plaques in different Alzheimer mouse models **[42]** as well as in human brain samples **[18,21]**. In this study, we were unable to detect any accumulation of GM1. Besides gangliosides, we detected certain phosphatidylinositol species with a similar accumulation pattern in APP/PS1 mouse brain. The feature *m/z* 559.4 putatively belonging to lysophosphatidylinositol (LPI) 18:0 and *m/z* 855.6 belonging to phosphatidylinositol 38:4 both show increased intensity in the ROIs defined by Aβ_1-42_ **(Figure 3b (v-vi))**. Whereas LPI 18:0 seemed to primarily appear in the vicinity of putative plaques, PI 38:4 was additionally located in the pyramidal layer of the hippocampus and to a certain extent in the cortex. Another lipid class that shows an altered behavior in the APP/PS1 mouse model are sulfoglycosphingolipids, especially sulfatides (ST). However, compared to the other lipid classes, they did not accumulate in the regions of interest. Rather, they appeared to be less intense in the immediate vicinity of the Aβ_1-42_-positive plaques, as compared to the control areas in the WT mouse **(Supplement Figure 2)**. We could detect depletion of two sulfatides (ST d42:2; and ST d42:1) in the APP/PS1 sample, especially in the cortex region at the Aβ1-42-positive regions. Besides this, their spatial distribution indicated that they were most intense in the white matter of both, the APP/PS1 and WT brain **(Figure 3b (vii-viii), Supplement Figure 2)**.

To investigate potential differences in the lipid composition of individual plaques, we focused on three ROIs. The white filled arrowhead pointing to an Aβ_1-42_-positive region in the cortex defines the first region **(Figure 3a (i))**. The same ROI in the lipid data set shows accumulation of gangliosides (i-iv), (lyso)-phosphatidylinositol (v-vi) as well as depletion of the two sulfatides (vii-viii). The white filled arrow indicates Aβ_1-42_-positive plaque ROI as well as only ganglioside and lyso-phosphatidylinositol (LPI 18:0) accumulation in the hippocampal region **(Figure 3b)**. The white unfilled arrows showing lipid deposition in the hippocampus which is only true for GM3 d36:1 and GM2 d36:1, but not for GM3 d38:1 and GM2 d38:1.

### 2.4. Structure elucidation of candidate lipid biomarkers by TOF/TOF on-tissue fragmentation using a bench top mass spectrometer

Our analysis demonstrates that not only for protein markers but also on the lipid level, differences in the microenvironment of individual Aβ plaque regions appear in the APP/PS1 mouse sample. To report accurate information of these potential lipid markers, a lipid TOF/TOF fragmentation analysis study was performed on the same benchtop MS to accurately identify the lipid species.

After acquisition of the MALDI MSI lipid data, we performed manual selection of precursors for on-tissue fragmentation to identify and annotate candidate lipid biomarkers described in the previous section.

Ganglioside GM2 d36:1 (18:1/18:0) (C_67_H_121_N_3_O_26_; *m/z* 1383.8 [M]) is an example of a candidate marker that we successfully fragmented **(Figure 4a)**. Chemical reference structures were obtained from LIPIDMAPS *[43]*. The corresponding fragmentation spectrum featured an intense peaks of the precursor *m/z* 1383.0 [M-H]^-^, suggesting that fragmentation efficiency could be further improved, if the MS^2^ was ambiguous. The peak at *m/z* 1091.8 corresponded to the diagnostic [GalGalGlcCer-H]^-^ ion without the sialic acid linkage. Finally, the fragment at *m/z* 290.0 displayed the counterpart, namely the N-acetylneuraminic acid linked to the inner Gal. The ion assignment was in accordance with published recommendations *[44]*. However, we could not detect any fatty acid fragments or the sphingolipid itself. Therefore, we cannot report on the exact arrangement and length of the respective fatty acyl chains **(Figure 4a bottom)**.

**Figure 4.**
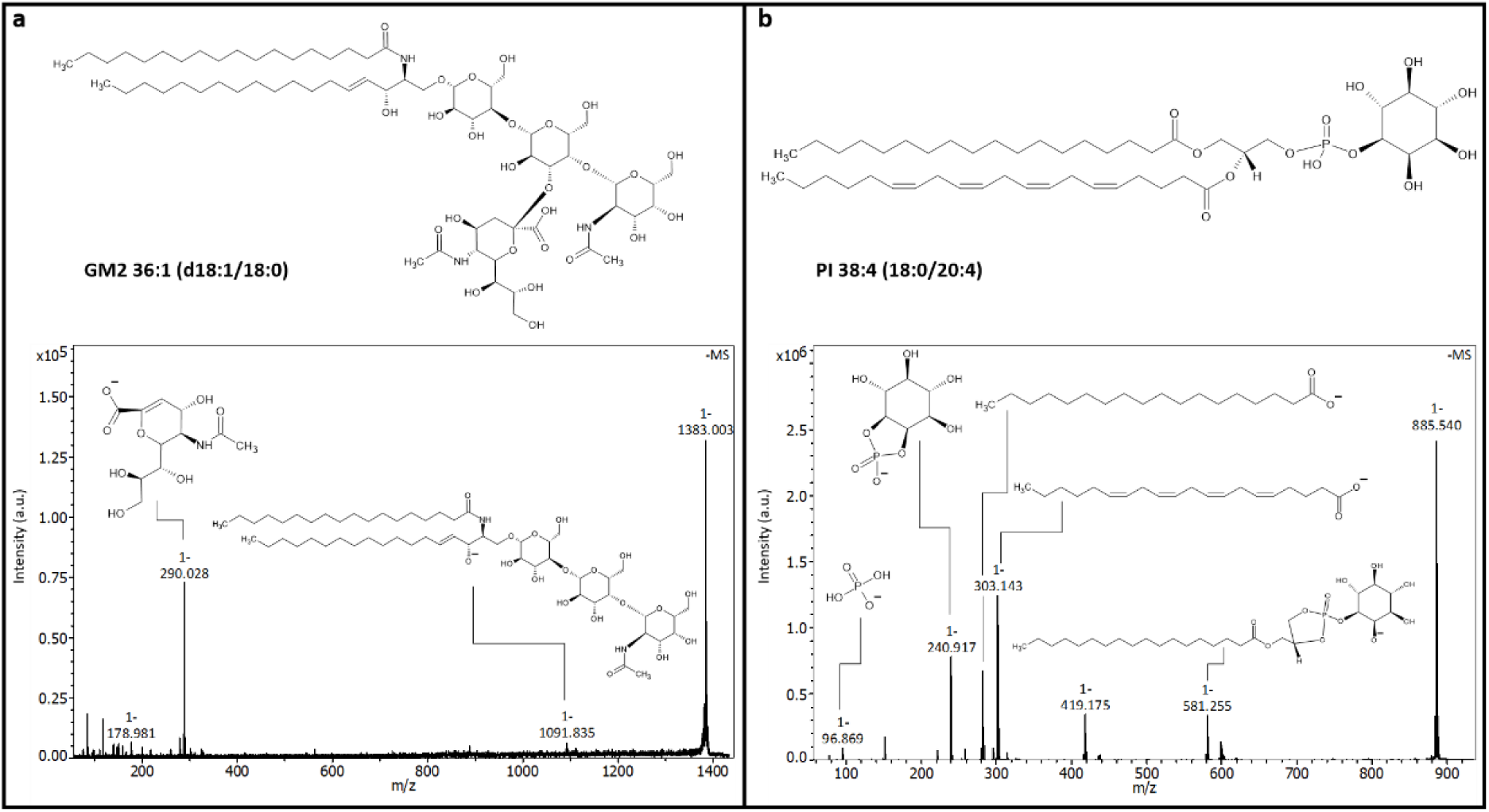
Identification of candidate lipid biomarkers by on-tissue TOF/TOF fragmentation analysis using a bench top mass spectrometer. **a,** Structure of ganglioside GM2 d36:1 (18:1/18:0) (C_67_H_121_N_3_O_26_; *m/z* 1383.8 [M]) andMS^2^ fragmentation spectrum of *m/z* 1383.0 [M-H]^-^ in negative ionization mode obtained on a neoflex TOF/TOF instrument. **b,** Structure of phosphatidylinositol 38:4 (18:0/20:4) (C_47_H_83_O_13_P; *m/z* 885.5 [M]) and MS^2^ fragmentation spectrum of *m/z* 885.5 [M-H]^-^ in negative ionization mode.

Similarly, the structure of phosphatidylinositol 38:4 (18:0/20:4) (C_47_H_83_O_13_P; *m/z* 886.5 [M]) was determined. The corresponding fragmentation spectrum, the precursor *m/z* 885.5 [M-H]^-^ was accompanied by a peak at *m/z* 581.3 corresponding to the triglyceride backbone connected to the inositol phosphate head group. Fragments *m/z* 303.1 and *m/z* 283.2 indicated the two fatty acid chains, whereas fragments *m/z* 241.0, *m/z* 153.0 and *m/z* 96.9 were indicative of the breakdown parts of the head group **(Figure 4b)**.

We performed on-tissue fragmentation of ten lipid *m/z* features in total and were able to successfully annotate all of them (Supplement Figure 3).

## Discussion

Multimodal analyses for pathological conditions such as AD are urgently needed to gain comprehensive insight into the various molecular and cellular processes of the disease (discussed in [45]). MSI offers the possibility to analyze a wide range of small molecules and reveal their spatial distribution within the tissue [14]. Combining this technique with HiPLEX-IHC enables the analysis of many relevant biomolecules from fatty acids to multi-kDa proteins. This study demonstrates that all measurements needed for such a multimodal analysis can be accomplished on a single bench top mass spectrometer. We successfully obtained data for lipids, crucial protein biomarkers, and histological information from a single tissue section derived from an AD model mouse. Our study revealed the co-localization of different protein biomarkers in the microenvironment of Aβ_1-42_-positive plaques in an APP/PS1 brain sample. We detected a high load of Aβ_1-42_-positive plaques in cortex and hippocampus and showed the accumulation of two important immune markers (Iba-1 and GFAP) at the plaque sites using photo-cleavable mass tag reporters. Additionally, we detected increased APP expression in the vicinity of plaques in the AD mouse model in comparison to the WT sample. Our findings suggest that co-localization between Aβ1_-42_ plaques and all three neurology markers is not absolute, but that there are noticeable differences in the molecular composition of the microenvironment of individual plaques. Upon further investigation of the lipid composition of Aβ_1-42_-positive plaques, distinct lipid classes were either accumulated or depleted in the ROI.

In this study, we established a method to detect Aβ plaques and activated microglia in the same tissue section. Microglia, the resident immune cells of the central nervous system, play a key role in brain homeostasis and neuroinflammation (reviewed in [46]). They have been observed in close association with amyloid-beta aggregates [47] and are now considered a potential third hallmark of AD alongside Aβ plaques and tau tangles [48]. In recorded ion images and overlays of the Aβ_1-42_ and Iba-1 signals we saw a high degree of co-localization of plaques and potentially clustered microglia in cortex and hippocampus, suggesting recruitment of microglia cells around amyloid beta plaques reported by others [49]. This observation is consistent with recent literature [50,51]. The question of whether plaque-associated microglia have a protective effect or contribute to progression of disease cannot be answered here. Earlier studies showed that microglia play a crucial role in AD, as they respond to Aβ material by attempting to clear the accumulations by phagocytosis or degrading enzymes [7,52]. Clustered microglia surround Aβ plaques are mainly responsible for phagocytosis of fibrillary Aβ peptides in APP/PS1 mice [50]. Thereby, microglia change from a resting into an reactive state that releases pro-inflammatory molecules to further drive neuro-inflammation [53].

Further support for enhanced neuro-inflammation from astrocytes in AD patient brains are also attracting increasing attention in AD research, as they are found as plaque components in AD animal models [54] and human AD patient samples [55]. GFAP is the main marker for astrogliosis, a process during which astrocytes transition into an reactive state during inflammation [56]. In this study, we identified GFAP-positive regions in the vicinity of Aβ_1-42_-positive plaques. This result is consistent with previous research the hippocampus of APP/PS1 model mice [6]. However, our results suggest that not all detected Aβ_1-42_ plaques have GFAP-positive astrocytes in their vicinity. This raises the question if an astrogliosis response might not be homogeneous throughout the affected brain areas. In different AD animal models, different results including complete lack of astrogliosis have been obtained by multiple laboratories. The absence of GFAP-positive astrocytes could indicate an impaired astrogliotic response, which could result from the failure of the immune system in the advanced course of plaque development (further discussed in the Review of [54].

Amyloid precursor protein (APP), a ubiquitously expressed transmembrane protein, plays a central role in AD as precursor of amyloid peptides like Aβ_1-42_. These Aβ peptides aggregate and form Aβ plaques [5]. In this study, we found accumulating APP signals mainly in cortex and hippocampus, mostly co-localizing with Aβ_1-42_-positive plaques. The APP/PS1 mouse model is genetically engineered to overexpress the human mutant APP, leading to increased amounts of APP protein in brains of these mice [10]. Therefore, the apparently clustered APP at the plaque site in the transgenic mouse could be an artifact of the AD model, which means that APP levels might not be increased or clustered to that extent at the plaque site in samples from patients with sporadic late-onset AD. Nevertheless, the Selkoe group tested different antibodies directed against APP in human AD patient samples and suggested the presence of full length APP in some but not every senile Aβ plaque [57]. In our experiments, APP staining revealed positive APP signal intensity in the pyramidal layers of CA and the granule layer of the dentate gyrus (DG). The Zheng group examined the localization of endogenous APP in mouse brain and found APP being specifically expressed in NeuN-positive neurons [58]. This result could explain our finding of increased APP-positive signal intensity in the APP/PS1 and WT sample in this exact region, which seems very similar to our NeuN signal distribution in the hippocampal area of the APP/PS1 and WT mouse sections.

In addition to protein markers, lipids have recently received increased attention in AD research. The transcriptomic signature of lipopolysaccharide (LPS)-activated microglia revealed changes in lipid- and lipoprotein-associated genes like ApoE and TREM2 [53], suggesting a strong link between microglial activation, lipid metabolism in the brain and potentially the neurodegenerative progression of the disease. Therefore, we investigated different lipid species using untargeted MSI.

In agreement with other animal and human studies, we detected GM, PI and ST species in close association with Aβ pathology [18,21,42]. Gangliosides represent a subclass of cell membrane glycosphingolipids. They are major components of lipid rafts in neuronal membranes and therefore essential for cell signaling processes and maintaining the activity of protein-protein and protein-lipid interaction [59,60]. In AD, there is increasing evidence of altered ganglioside metabolism [9]. We detected increased levels of GM2 and GM3 species in the APP/PS1 mouse model in cortex and hippocampus, thus confirming earlier MSI studies [18,42]. As in some published mouse studies, we did not detect significant accumulations of GM1 in the surrounding of Aβ1-42-positive plaques, [23,42]. In healthy brains, complex gangliosides like GM1 are the predominant form, whereas the simpler forms (GM2, GM3) may be expressed in small quantities [60]. Low levels of GM1 and increased levels of GM2 and GM3 could indicate an altered lysosomal degradation of gangliosides. Potentially, GM1 is degraded more than the other gangliosides, resulting in more accumulation of GM2 and GM3 [42]. However, the exact role of gangliosides in AD progression is not yet fully understood and needs further investigation. Besides gangliosides, we observed increased phosphatidylinositols at plaque sites in APP/PS1 brain. Altered phosphatidylinositol levels have been already reported in association with other AD mouse models and human AD samples [20,21]. Interestingly, the observed PI (18:0/20:4) species yields the observed lysophosphoinositol (LPI 18:0) and arachidonic acid (AA) upon phospholipase A2 cleavage. Arachidonic acid (AA) is the precursor in eicosanoid biosynthesis and is known to be involved in microglial and astroglial inflammatory response mechanisms [61]. Phospholipase A2 (PLA2) was previously found to be upregulated in AD brains [62]. Higher mRNA expression was found to be connected to astrocytes associated with Aβ plaques in AD brain [63]. Additionally, lysophospholipids (LPIs), the degradation products of phospholipids, are produced by cytosolic phospholipase A2 (cPLA2). These LPIs seem to be increasingly released by neurons and are capable of inducing astrogliosis and therefore regulating neuroinflammation in AD [64].

Besides accumulation patterns of different lipid species, we detected one lipid class (sulfatides) that were depleted at plaque sites. Sulfatides are important lipids in the central nervous system and are major components of the myelin sheath [65], with MSI studies already reporting the depletion of sulfatides at amyloid plaque sites [42]. An LCMS study with human AD patient samples confirmed this finding and showed a decrease in sulfatides in this samples as well as noticeable increase in ceramide content, which could be the sulfatide degradation product [66].

In conclusion, this study showed the level of molecular complexity in AD tissue sample that can be revealed by our workflow on a single tissue section with a benchtop mass spectrometer. Besides Aβ peptides, many different molecules are present in the plaque microenvironment, with each of them potentially playing an important role in disease progression. Therefore, it is not enough to consider each molecule individually, but within the context of the tissue environment. Only a thorough understanding of the interaction between proteins, small molecules, and different cell types will promote a more refined understanding of AD pathogenesis and reveal new drug discovery avenues. In future studies of the metabolism of Aβ plaques, this multiomic and multimodal method may impact and be replicated with other mouse models or human AD patient samples.

## 4. Materials and Methods

### Chemicals

All chemicals and solvents were of high-performance liquid-chromatography mass spectrometry (HPLC-MS) grade. Acetonitrile (ACN), acetone, water, ethanol (EtOH), sodium chloride, acetone, and 2-propanol were from VWR Chemicals (Darmstadt, Germany). Trifluoroacetic acid (TFA), α-cyano-4-hydroxycinnamic acid (α-CHCA), octyl β-D-glycopyranoside (OBG), phosphate buffered saline (PBS), alkaline retrieval buffer, chloroform, paraformaldehyde, hydrochloric acid and the hydrophobic barrier pen were from Merck KGaA (Darmstadt, Germany). Vacuum-stable caged MALDI matrix 4,5-dimethoxy-2-nitrobenzyl-2,5-dihydroxyacetophenone (DMNB-2,5-DHAP) [38] was provided by Sirius Fine Chemicals GmbH (SiChem, Bremen, Germany). Rabbit serum (sterile filtered) was purchased from Capricorn Scientific (Ebsdorfergrund, Germany), sodium hydroxide from Fisher Scientific (Hampton, NH, USA), and Tris-HCl and Tris base were purchased from Carl Roth GmbH (Karlsruhe, Germany). HiPLEX Miralys “Neurology Panel” antibodies labeled with photo-cleavable mass-tags (PCMT) were obtained from AmberGen Inc. (Billerica, MA, USA).

### Animal and Tissue collection

APP PS1-21 and the non-transgenic littermate (C57BL/6J) mice [10] were obtained from the Jucker lab and bred for Abbvie by Charles River Laboratories (Sulzfeld, Germany). The mice were kept in a temperature- and humidity-controlled room with a 12:12 hour dark/light cycle with ad libitum access to water and food. All animal experiments were performed in full compliance with the Principles of Laboratory Animal Care [67] in an AAALAC accredited program where veterinary care and oversight was provided to ensure appropriate animal care. All animal studies were approved by the government of Rhineland Palatinate (Landesuntersuchungsamt) and conducted in accordance with the directive 2010/63/EU of the European Parliament and of the Council on the protection of animals used for scientific purpose, the ordinance on the protection of animals used for experimental or scientific purposes (German implementation of EU directive 2010/63; BGBl. I S. 3125, 3126), the Commission Recommendation 2007/526/EC on guidelines for the accommodation and care of animals used for experimental and other scientific purposes, the German Animal Welfare Act (BGBl. I S. 1206, 1313) amended by Art. 1 G from 17 December 2018 I 2586.

Fresh-frozen brains were cut into 10 *µ*m thick coronal sections at −15°C using a Leica CM 1860 UV cryostat (Leica Biosystems, Nussloch, Germany). Tissue sections were thaw-mounted onto conductive MALDI IntelliSlides™ (Part No 1868957; Bruker Daltonics GmbH, Bremen, Germany) and stored in vacuum bags at −80°C. Prior to MS analysis, tissue sections were thawed under vacuum for approximately 1 hr.

### Sample preparation MALDI Mass Spectrometry Imaging

For lipid MSI, tissue was spray-coated with 2.5 mg/mL DMNB-2,5-DHAP matrix [38] in 80% ACN, 20% water and 0.1 % TFA using a TM sprayer (HTX Technologies, Chapel Hill, NC, USA) with the following spray parameters: flow rate of 0.1 mL/min, spray temperature 75°C, nozzle height at 40 mm, ten layers sprayed in an HH pattern, with spray velocity of 1200 mm/min.

Prior to MALDI HiPLEX-IHC staining, MALDI matrix was removed with 2x 3 min washes in ice-cold acetone at -80°C and then dried under vacuum for 10 min. This was followed by the following steps in glass Coplin jars at room temperature (RT): Tissue sections were fixed in freshly prepared 1% paraformaldehyde (PFA) for 30 min followed by a 10 min wash in PBS. Lipids and metabolites were then removed in two 3 min acetone washes followed by 3 min in Carnoy’s solution. Next, the sections were rehydrated using a series of water/EtOH washes, i.e., 2x for 2 min in 100% EtOH, 3 min in 95% EtOH, 3 min in 70% EtOH and 3 min in 50% EtOH. After rehydration, sections were equilibrated in 1X Tris-buffered saline (TBS) (10X stock: 50 mM Tris, 2 M NaCl, pH 7.5) for 10 min. Antigen retrieval was performed for 30 min in alkaline Tris-EDTA buffer pH 9 in a water bath at 95°C, before being equilibrated at RT for 30 min. After antigen retrieval, the sections were again washed for 10 min in TBS and blocked in tissue blocking buffer (TBB: 2% Serum, 5% BSA in TBS with 0.05% OBG) for 1 hr. PCMT-labeled antibodies were diluted in TBB and filtered (0.45 *µ*m micro-centrifuge filter unit) for 1 min. Each tissue section on the slide was circled with a hydrophobic pen to retain small fluid volumes, and the slide was then placed in a humidity chamber. All following steps were performed in the dark. 70 *µ*L antibody probes were applied per section, and the slide was incubated overnight at 4°C. Slides were removed from the humidity chamber and washed 3x for 5 min in TBS with gentle shaking. Next, sections were rebuffered for 10 s in ammonium bicarbonate (ABC) buffer, followed by 3x for 2 min in ABC buffer with gentle shaking. After removal of excess fluid, sections were vacuum-dried for 1.5 hr in a desiccator. Mass tags were photo-cleaved for 10 min using a near-UV Light Box (Ambergen). Next, 10 mg/mL α-CHCA in 70% ACN and 0.1% TFA were spray-coated in six passes at 60°C, a flow rate of 0.07 mL/min with a velocity of 1200 mm/min, a track spacing of 2 mm at 10 psi pressure and a drying time of 10 s in a crisscross pattern using a HTX M5 sprayer (HTX Technologies). For re-crystallization of the HCCA matrix, the slide was incubated in a petri dish with a filter paper (S&S, Schleicher & Schüll, Germany) soaked in 1 mL 5% isopropanol at 55°C for 1 min. If not imaged immediately, the slide was stored at 4°C in a vacuum-sealed slide mailer.

### MALDI MS Imaging (MSI) and MS/MS Data Acquisition

Lipid MSI (*m/z* 400−1600; 200 shots per pixel) of coronal mouse brain sections was performed in reflector negative ionization mode at 20 *μ*m pixel size using a neofleX benchtop mass spectrometer (Bruker Daltonics, Bremen, Germany) equipped with a 10 kHz laser. External cubic-enhanced calibration was performed using red phosphorous spotted adjacent to the tissue sections. For tandem-MS analysis, relevant lipids in the amyloid-β plaque region were fragmented using the TOF/TOF capability of the neofleX, with a laser ablation area of 50x50 *µ*m. Precursor masses were selected for fragmentation with an isolation window of 0.65 % of the precursor mass.

MALDI HiPLEX-IHC MSI (*m/z* 600-4000) was conducted in reflector positive ionization mode at 20 µm pixel size with a sampling rate of 1 Gs/s. Matrix suppression was set up to 640 *m/z*. For data processing, the smoothing algorithm was Savitzky-Golay, with a width of 0.01 *m/z* and 5 cycles. Centroid peak detection was used with S/N threshold of 6, a peak width of 0.05 *m/z* and peak height of 80%. Baseline subtraction was performed with the TopHat algorithm.

### Hematoxilin & eosin (H&E) staining

For H&E staining after lipid MSI and MALDI HiPLEX-IHC analysis, the MALDI matrix was removed by 1 min incubation in 70% ice-cold EtOH. Slides were placed in hematoxylin (Merck) for 3 min, washed with Milli-Q water for 1 min, dipped into acidic alcohol (70% EtOH with 0.1% HCl), and then washed again with Miili-Q water for 1 min. To increase the pH, a blueing step was performed (Blue solution stock: 10 g Na_4_HCO_3_ + 100 g magnesium sulfate in 1 L of Milli-Q water) for 2 min, and then slides were rinsed with Milli-Q water for 1 min. Then the slide was counterstained in 0.5% eosin (Merck) for 2 min and washed in Milli-Q water for 1 min. The section was finally washed and dehydrated in 80% EtOH, 96% EtOH, and 100% EtOH for 1 min each. Finally, the tissue section was dipped into xylene (Merck) for a few seconds and then mounted with Eukitt (Merck), before microscopic images were acquired using an AperioCS2 scanner (Leica Biosystems).

### Data analysis

MALDI HiPLEX-IHC data was visualized in SCiLS Scope 1.0 software (Bruker Daltonics). Individual channels for all PCMTs were displayed and evaluated (Table 1). Image registration of MALDI Lipid MSI and MALDI HiPLEX-IHC was performed using in-house M^2^aia software [39,40]. Raw spectra were imported into SCiLS Lab (Bruker Daltonics).

**Table 1.** Theoretical masses of photo-cleavable and mass-tagged (PCMT) antibodies, their corresponding antigens and concentrations.

| Antigen | PCMT theoretical mass | Concentration [µg/mL] |
| --- | --- | --- |
| Iba-1 | 959.57 | 3.75 |
| Amyloid-β<sub>1-42</sub> (Aβ<sub>1-42</sub>) | 1770.87 | 3.75<sup>1</sup> |
| GFAP | 1011.54 | 3.75 |
| NeuN | 1308.70 | 3.75 |
| APP | 1722.93 | 5.00 |

## Author Contributions

Conceptualization, E.M., K.B., C.H.; methodology, E.M., T.E., D.N., L.v.A., C.H.; software/data science: T.E.; validation, E.M., T.E., D.N., K.B.; investigation, E.M., T.E., L.v.A., D.N., K.B.; resources, C.K., C.H.; data curation, E.M., T.E., D.N.; writing—original draft preparation, E.M., K.S., C.H.; writing—review and editing, all; visualization, E.M., T.E., D.N.; supervision, X.X.; project administration, E.M., K.B., C.H.; funding acquisition, K.B., C.K., C.H. All authors have read and agreed to the published version of the manuscript.”

## Institutional Review Board Statement

All animal experiments were performed in full compliance with the Principles of Laboratory Animal Care [67] in an AAALAC accredited program where veterinary care and oversight was provided to ensure appropriate animal care. All animal studies were approved by the government of Rhineland Palatinate (Landesuntersuchungsamt) and conducted in accordance with the directive 2010/63/EU of the European Parliament and of the Council on the protection of animals used for scientific purpose, the ordinance on the protection of animals used for experimental or scientific purposes (German implementation of EU directive 2010/63; BGBl. I S. 3125, 3126), the Commission Recommendation 2007/526/EC on guidelines for the accommodation and care of animals used for experimental and other scientific purposes, the German Animal Welfare Act (BGBl. I S. 1206, 1313) amended by Art. 1 G from 17 December 2018 I 2586.

## Data Availability Statement

Data will be available from the corresponding author after publication upon reasonable request.

## Acknowledgements

We are grateful to Lars Gruber for sharing his expertise in MS/MS data analysis and his support in setting up the neoflex instrument in our lab. We thank Tobias Bausbacher for establishing the automated H&E staining protocol. We also acknowledge Jonas Cordes for the introduction to the software M2aia. Finally, we thank Stefan Schmidt for insightful discussions and reviewing the figures.

