## Supplementary Data for "Exploring the Aβ Plaque Microenvironment in Alzheimer’s Disease Mice by Multimodal Lipid-Protein-Histology Imaging on a Benchtop Mass Spectrometer"

**Supplementary Figures**

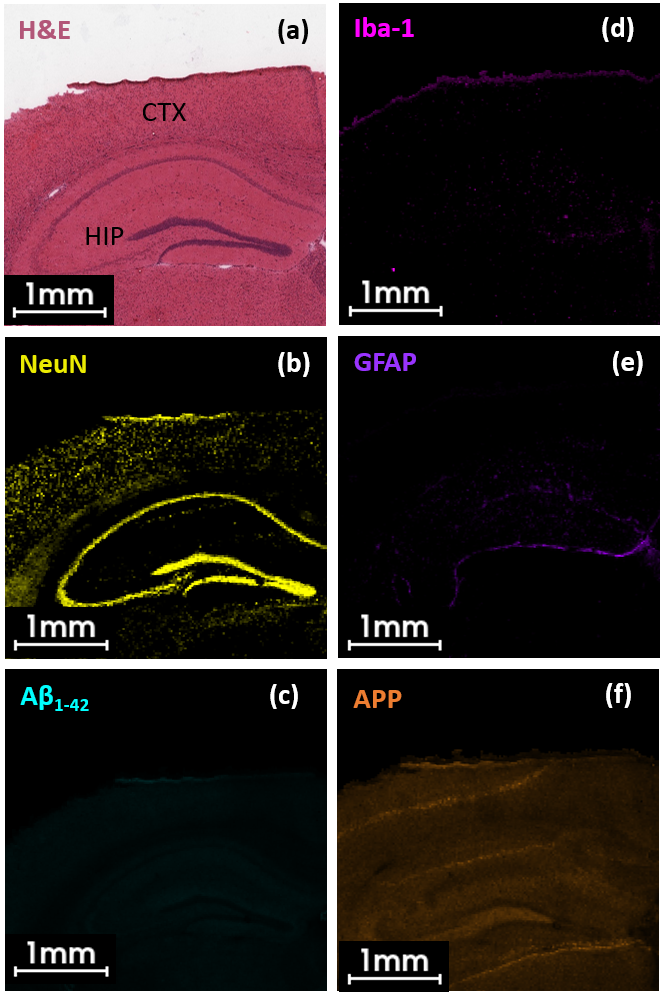

Supplementary Figure 1. Endogenous protein biomarker distribution in WT mouse brain visualized by MALDI-HiPLEX IHC

(a) H&E staining highlight distinct brain structures e.g., the hippocampus and cortex (CTX). HiPLEX-MALDI IHC images of five PCMTs visualize marker distribution in a coronal WT mouse brain section. Single ion images of brain structural marker(b) NeuN (Neuronal nuclei, m/z 1309.0 [M+H]^+^), (c) AD amyloid-β _1-42_ peptide (Aβ_1-42_, m/z 1771.5 [M+H]^+^), (d) microglia marker Iba-1 (m/z 960.1 [M+H]^+^), (e) glial fibrillary acidic protein (GFAP, m/z 1011.9 [M+H]^+^) and (f) amyloid precursor protein (APP, m/z 1723.6 [M+H]^+^).

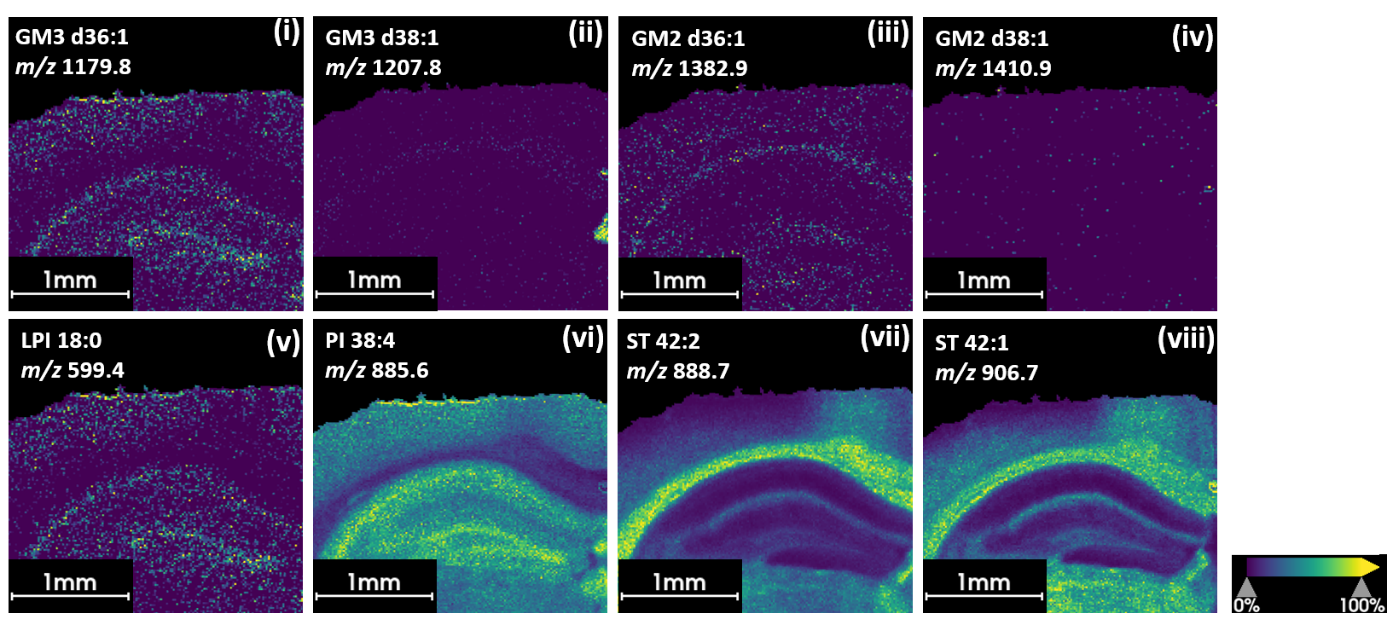

Supplementary Figure 2. MALDI lipid imaging reveals distribution of selected lipids in WT mouse brain tissue

**a,** MALDI lipid ion images of eight different lipid features in negative ionization mode detected in WT mouse brain **(i-iv)**. Ganglioside GM3 d36:1 (m/z 1179.8, [M-H]^-^), GM3 d38:1 (m/z 1207.8, [M-H]^-^), GM2 d36:1 (m/z 1382.9, [M-H]^-^), GM2 d38:1 (m/z 1410.9, [M-H]^-^), lyso-phosphatidylinositol 18:1 (LPI, m/z 599.4, [M-H]^-^), phosphatidylinositol 38:4 (PI m/z 885.6, [M-H]^-^), sulfatide 42:2 (ST d42:2;O2, m/z 888.7, [M-H]^-^) and sulfatide 42:1 (ST d41:2;O2, m/z 906.7, [M-H]^-^).

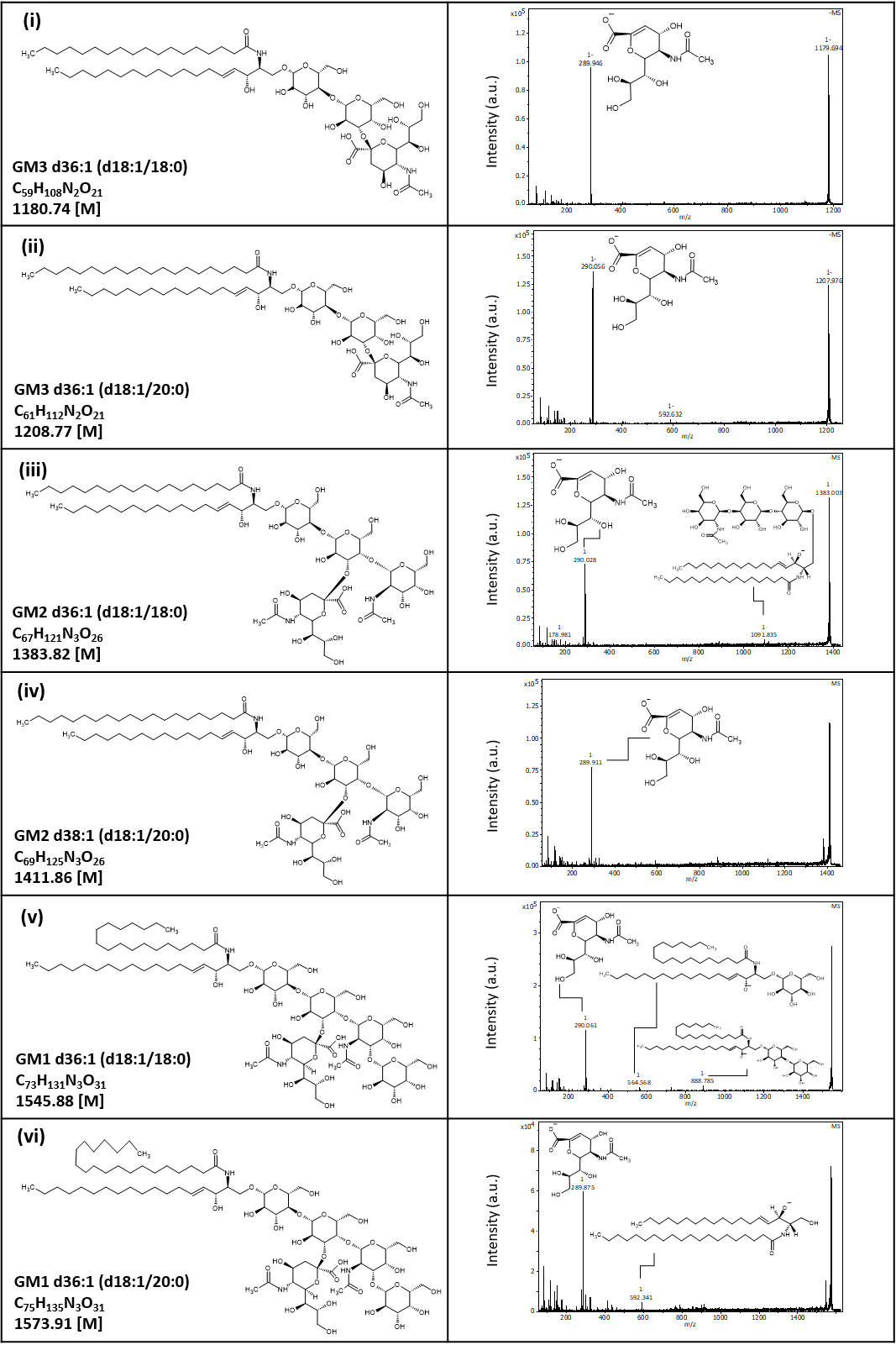

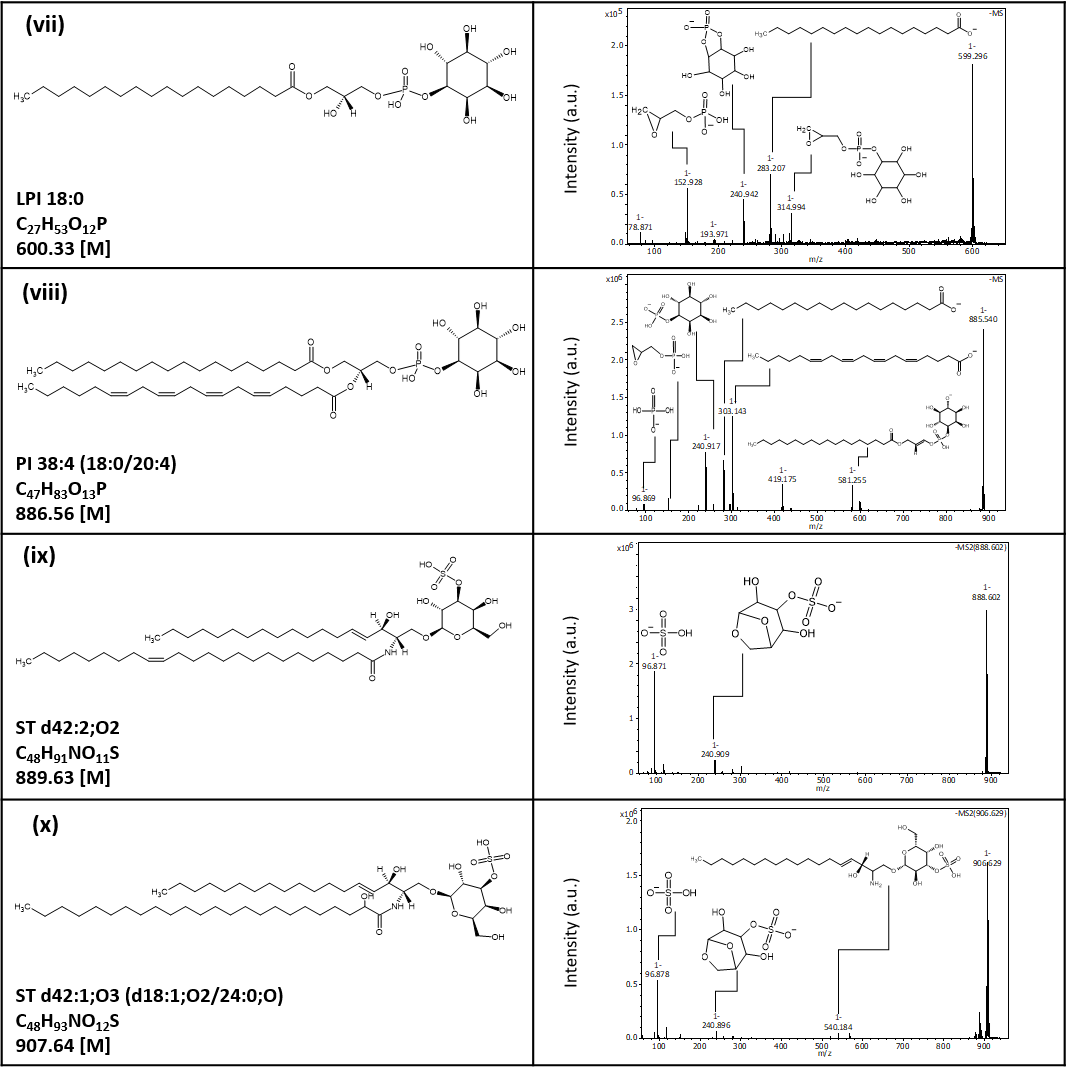

Supplementary Figure 3. Identification of candidate AD lipid biomarkers by TOF/TOF on-tissue fragmentation analysis using a bench top mass spectrometer

Summary of the chemical structure of the putative lipid precursor of interest (left) and the respective MS^2^ fragmentation spectra from single shot data acquisition with identified lipid fragments (right) obtained on a neoflex MALDI-TOF/TOF benchtop instrument (i-x).

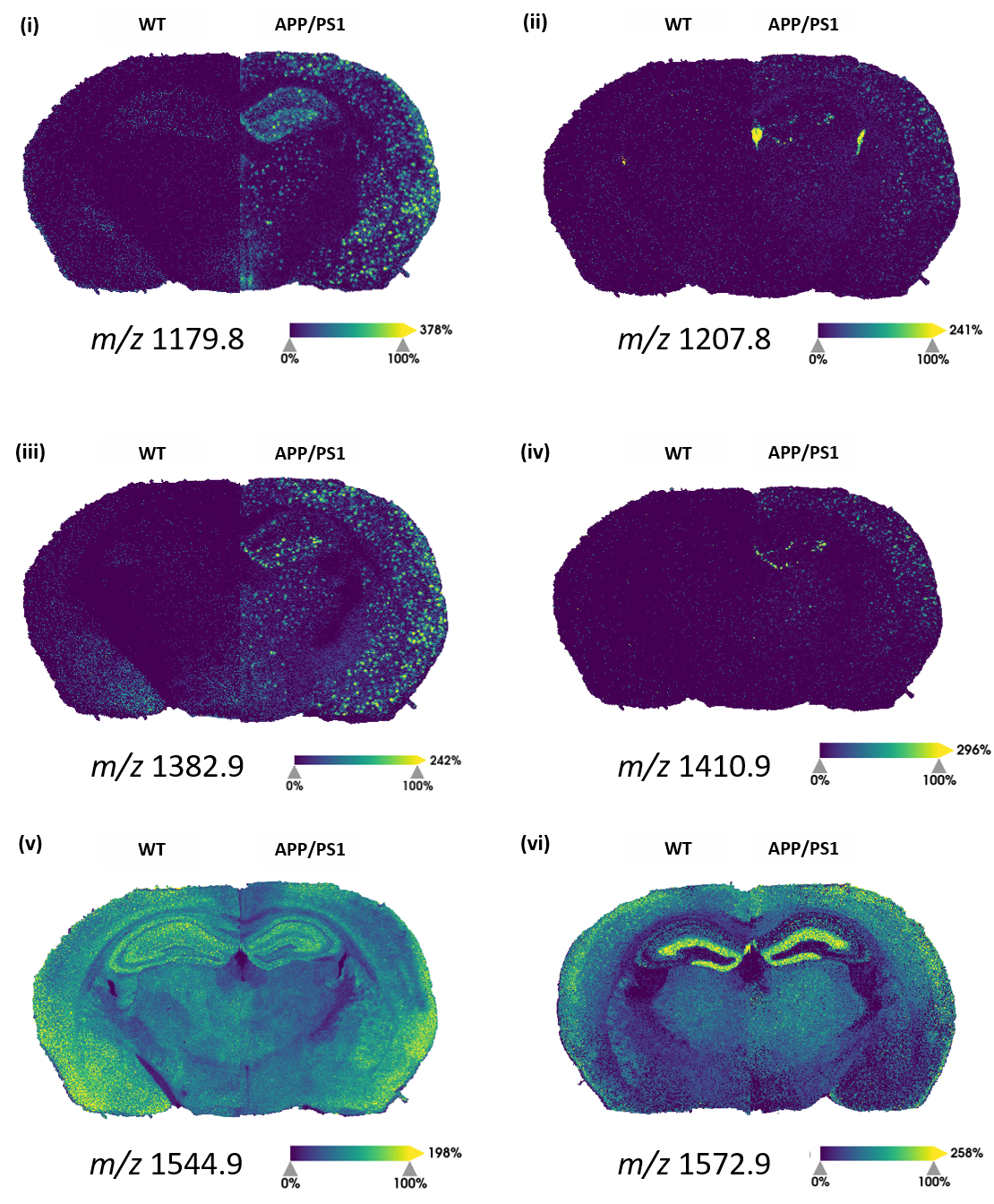

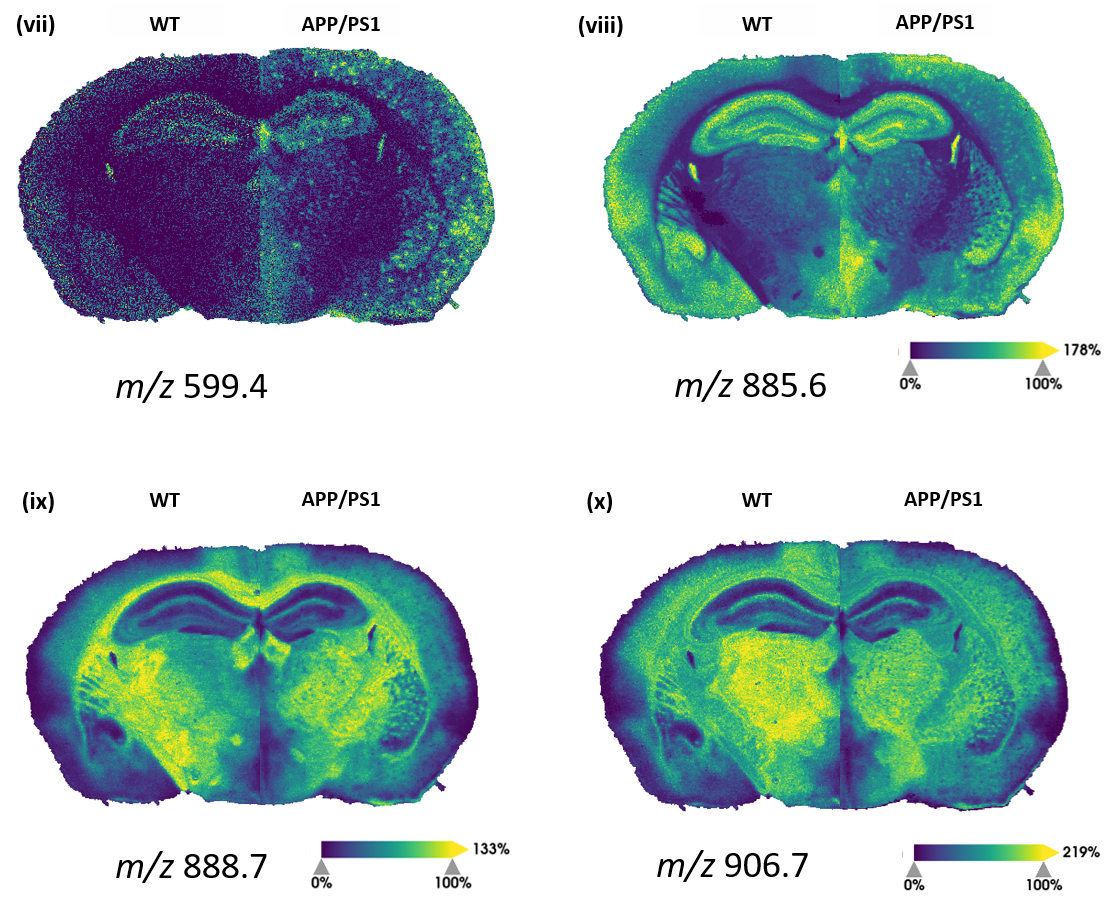

Supplementary Figure 4. Single MS1 ion images of ten lipids of interest in APP/PS1 and wild-type (WT) mouse brain tissue.

Summary of single ion images of ten different lipids obtained on a neoflex MALDI-TOF/TOF benchtop mass spectrometer (i-x). Ion images show half a mouse brain of the WT mouse and half a brain of the APP/PS1 mouse.
